# PLaBAse: A comprehensive web resource for analyzing the plant growth-promoting potential of plant-associated bacteria

**DOI:** 10.1101/2021.12.13.472471

**Authors:** S. Patz, A. Gautam, M. Becker, S. Ruppel, P. Rodríguez-Palenzuela, DH. Huson

**Affiliations:** Institute for Bioinformatics and Medical Informatics, University of Tuebingen, Tuebingen, Germany; International Max Planck Research School “From Molecules to Organisms”, Max Planck Institute for Developmental Biology, Tuebingen, Germany; Institute for National and International Plant Health, Julius Kuehn-Institute - Federal Research Centre for Cultivated Plants, Braunschweig, Germany; Leibniz Institute of Vegetable and Ornamental Crops, Grossbeeren, Germany; Centre for Plant Biotechnology and Genomics, Universidad Politécnica de Madrid-Instituto Nacional de Investigación y Tecnología Agraria y Alimentaria, Madrid, Spain

**Keywords:** plant-associated bacteria, plant growth-promotion, ontology, database, inoculant, plant production, agriculture

## Abstract

Plant-beneficial microorganisms are gaining importance for sustainable plant production and phytosanitary practices. Yet there is a lack of computational approaches targeting bacterial traits associated with plant growth-promotion (PGP), which hinders the *in-silico* identification, comparison, and selection of phytostimulatory bacterial strains. To address this problem, we have developed the new web resource PLaBAse (v1.01, http://plabase.informatik.uni-tuebingen.de/pb/plabase.php), which provides a number of services, including (i) a database for screening 5,565 plant-associated bacteria (PLaBA-db), (ii) a tool for predicting plant growth-promoting traits (PGPTs) of single bacterial genomes (PGPT-Pred), and (iii) a tool for the prediction of bacterial plant-association by marker gene identification (PIFAR-Pred). The latter was developed by Martínez-García et al. and is now hosted at University of Tuebingen. The PGPT-Pred tool is based on our new PGPT ontology, a literature- and OMICs-curated, comprehensive, and hierarchical collection of ∼6,900 PGPTs that are associated with 6,965,955 protein sequences. To study the distribution of the PGPTs across different environments, we applied it to 70,540 bacterial strains associated with (i) seven different environments (including plants), (iii) five different plant spheres (organs), and (iii) two bacteria-induced plant phenotypes. This analysis revealed that plant-symbiotic bacteria generally have a larger genome size and a higher count of PGPT-annotated protein encoding genes. Obviously, not all reported PGPTs are restricted to -or only enriched in-plant-associated and plant symbiotic bacteria. Some also occur in human- and animal-associated bacteria, perhaps due to the transmission of PGP bacteria (PGPBs) between environments, or because some functions are involved in adaption processes to various environments. Here we provide an easy-to-use approach for screening of PGPTs in bacterial genomes across various phyla and isolation sites, using PLaBA-db, and for standardized annotation, using PGPT-Pred. We believe that this resource will improve our understanding about the entire PGP processes and facilitate the prediction of PGPB as bio-inoculants and for biosafety strategies, so as to help to establish sustainable and targeted bacteria-incorporated plant production systems in the future.

## INTRODUCTION

A changing climate causing increasingly extreme weather events calls for novel strategies to improve crop resilience in order to maintain and guarantee global nutrition in the future, according to the WHO ^1^. In the past decades, plant-centric approaches, such as crop-plant breeding and the use of genetically modified organisms (GMOs), have received much attention. Additionally, new insights into the natural phytobiome have boosted the interest in naturally occurring phytostimulatory plant- and plant-soil-associated bacteria ^2–5^. Already in 1947, Allison et al. and other researchers thereafter, postulated the use of symbiotic microbial plant inhabitants to enhance growth and vitality of crop plants under different environmental conditions ^6–8^. Nowadays, the genetic and metabolic activity of plant growth-promoting bacteria (PGPBs) and fungi (PGPFs) are studied to evaluate their potential as agricultural bio-inoculants ^9,10^. Known efficient bacterial bio-inoculant strains, aside from *Rhizobiaceae* (comprising diazotrophic bacteria associated with root nodules of legumes), are limited to a few species from a low number of genera, though scattered across the domain of bacteria, like *Azospirillum* spp. (e.g., Nodumax-L), *Bacillus* spp. (e.g., RhizoVital®) or *Bradyrhizobium* spp. (e.g., applied to soybeans in Argentina) ^11–13^. Currently, other promising strains come to light, whose efficiency to promote plant biomass production and yield, even under various environmental stress conditions, has repeatedly been confirmed, such as for species of *Kosakonia, Pseudomonas*, and *Cronobacter*, albeit some of the PGPB-comprising genera also share pathogenic strains ^14–18^. Studies have revealed a huge spectra of microbial plant-beneficial properties, such as plant protection against pathogens, resistance to extreme temperature, bioremediation or early flowering ^19–21^. However, soil quality and management, plant genotypes and environmental conditions have a significant impact on the plant-beneficial microbial community ^22,23^. Hence, even if plant growth-promoting effects of up to 30% and more have been reported, varying environmental conditions and fragmentary understanding of genetic and metabolic microbial site-adaption processes have often caused a lower beneficial impact of such inoculants than expected.

Continuous improvement of OMICs techniques and bioinformatics algorithms for post-analysis contribute to a better understanding of the composition of host-associated microbiomes and of the host-microbiome interactions. Online resources, like the “CIRM-Plant-associated Bacteria/CIRM-CFBP” (https://www6.inrae.fr/cirm_eng/CFBP-Plant-Associated-Bacteria) database, contain up to ∼6,500 plant-associated bacteria and food-borne pathogens, affiliated with 60 genera ^24,25^. While a large number of resources, such as eggNOG ^26^, GO (Gene Ontology) ^27–29^, InterPro ^30^, KEGG ^31,32^, SEED ^33,34^, Bio/MetaCyc ^35,36^ and CAZy ^37^ facilitate general functional annotation, the plant-specific genetic repertoire of these strains has still to be elucidated. Recent bioinformatics approaches aim to reveal novel genetic and metabolic elements involved in plant-bacteria interaction ^38,39^. In 2016, Mwita et al. gave a deeper insight into active genetic elements and gene regulation of plant-associated microbes in response to root exudates by transcriptome analyses ^40^. In 2018, Levy et al. surveyed orthologous groups of bacterial proteins and domain profiles that appear to be characteristic of plant-vs. non-plant- and root-vs. soil-associated bacterial strains or microbial communities ^41^. In 2015, Bulgarella et al. provided evidence for a positive selection of genetic traits that contributes to a stabile host-microbe interaction ^42^. In 2016, Martínez-García et al. developed a supervised machine-learning approach to facilitate the prediction of plant-associated bacteria based on ∼170 manually curated traits, applicable as the PIFAR tool ^43^. More recently, Carrión et al. provided a detailed overview of largely unknown disease-suppressive plant-microbial features. In that approach metagenome-assembled genomes (MAGs) were analyzed regarding biosynthetic gene clusters (BGCs) using dbScan ^44^, AntiSMASH ^45–49^ and BIG-SCAPE ^50^ and further compared to the MIBiG database ^51,52^. In 2020, the proGenomes2 database was announced, providing accurate host associations and functional classifications of prokaryotic genomes, including plant-associated bacteria.

None of these approaches explicitly summarizes the plant symbiotic and growth-promoting genetic traits (PGPTs) as one entity ^53^, although a number of different plant-endophytic genetic and metabolic processes, important for a symbiotic lifestyle (e.g., host colonization, host interaction and host vigor enhancement), have been studied ^54,55^.

Glick (2012), Gouda et al. (2018) and Afzal et al. (2019), among others, provide overviews of functional plant growth-promoting processes and divide them on a first level into direct and indirect plant-beneficial effects ^56–58^. While direct effects comprise mechanisms involved in bio-fertilization, bioremediation and plant growth regulation, indirect effects encompass mechanisms responsible for plant biotic and abiotic stress control (e.g., biocontrol), induction of plant systemic resistance (ISR), induction of systemic acquired resistances (SAR), microbial host colonization and competitive exclusion ^59–66^. Although considerable efforts have been made to unveil the nature of plant-microbe association and of unknown biochemical compounds, the majority of these results are usually restricted to only a few taxonomic groups, lack biological meaningful explanations for respective genes or lack a clear separation between phytopathogenic and plant-beneficial strains. Similarly, publications that aim at describing entire PGPTs within a single bacterial genome often miss important features, for example, the entire vitamin biosynthesis or traits of competitive exclusion and fitness (e.g., toxin-antitoxin systems), which hampers comparability. Keep in mind that mechanisms for plant growth-promotion, especially related to indirect effects, may be essential for invading other environments and thus, also for pathogens.

Our study addresses the issues related to the prediction of PGPTs and comparability of annotation results. Here, we set up a webpage for plant-associated bacteria, called PLaBAse (http://plabase.informatik.uni-tuebingen.de/pb/plabase.php), where our hierarchical plant growth-promoting traits (PGPT) ontology is connected to plant-associated bacteria that are stored in PLaBA-db and implemented in the PGPT-Pred web-tool. Based on this, PGPT-Pred facilitates standardized annotation of PGPTs for (i) known PGPBs, (ii) not yet classified putative PGPBs, including those of other environments than plant, and (iii) pathogenic strains. We applied our ontology to ∼72,000 bacterial genomes stored on the JGI / IMG Genome Server, to demonstrate the potential of PLaBAse for studying the distribution of PGPTs, not only in the context of plant-associated (PA) bacteria and plant symbionts (e.g., PGPBs), but also regarding their abundancies in other environments (e.g., human, animal, aquatic, industrial).

## MATERIAL AND METHODS

### Web Server setup for PLaBAse

“PLaBAse” (http://plabase.informatik.uni-tuebingen.de/pb/plabase.php) is a linux-based web server (see Figure 1) that (i) provides access to the plant-associated bacteria database PLaBA-db (http://plabase.informatik.uni-tuebingen.de/pb/plaba_db.php), that (ii) carries out fast annotation of PGPTs via the PGPT-Pred tool (v1.1.0, http://plabase.informatik.uni-tuebingen.de/pb/form.php?var=PGPT-Pred), and (iii) analyzes plant interaction and virulence factors via the PIFAR-Pred tool (v1.2.0, http://plabase.informatik.uni-tuebingen.de/pb/form.php?var=PIFAR-Pred). The web interface is based on HTML5 and bootstrap. The backend pipeline uses PHP (v7.2.24), Perl (v5.26.1), Python3 (v3.7.4), and R (v4.0.2). Further, we have implemented a queuing system for handling job submissions using PHP and MySQL (v5.7.34). Users are notified through email on submission and completion of their job(s), or in case of any issues. By default, the job submission form for either tool executes the blastp+hmmer approach (v2.10.1+, v3.3) as described by Martínez-García et al ^43^. After submission, all genomic protein sequences are aligned against the tool-specific protein database and best hits are selected using an e-value threshold of 1e-5. The presence of respective PFAM domains is determined using hmmer, as described by Martínez-García et al ^43^. Additionally, we have included the Python-based IMG-KEGG-annotation_MAPPER in the PGPT-Pred tool, which enables mapping of KEGG annotations of the IMG database against our PGPT ontology, described below. The results page displays all data as online plots generated by Plotly (v1.58.5) including the open-source JavaScript graphing library, and KronaTools (v2.8)^67^, and allows download of results as tab-separated files.

**Figure 1:**
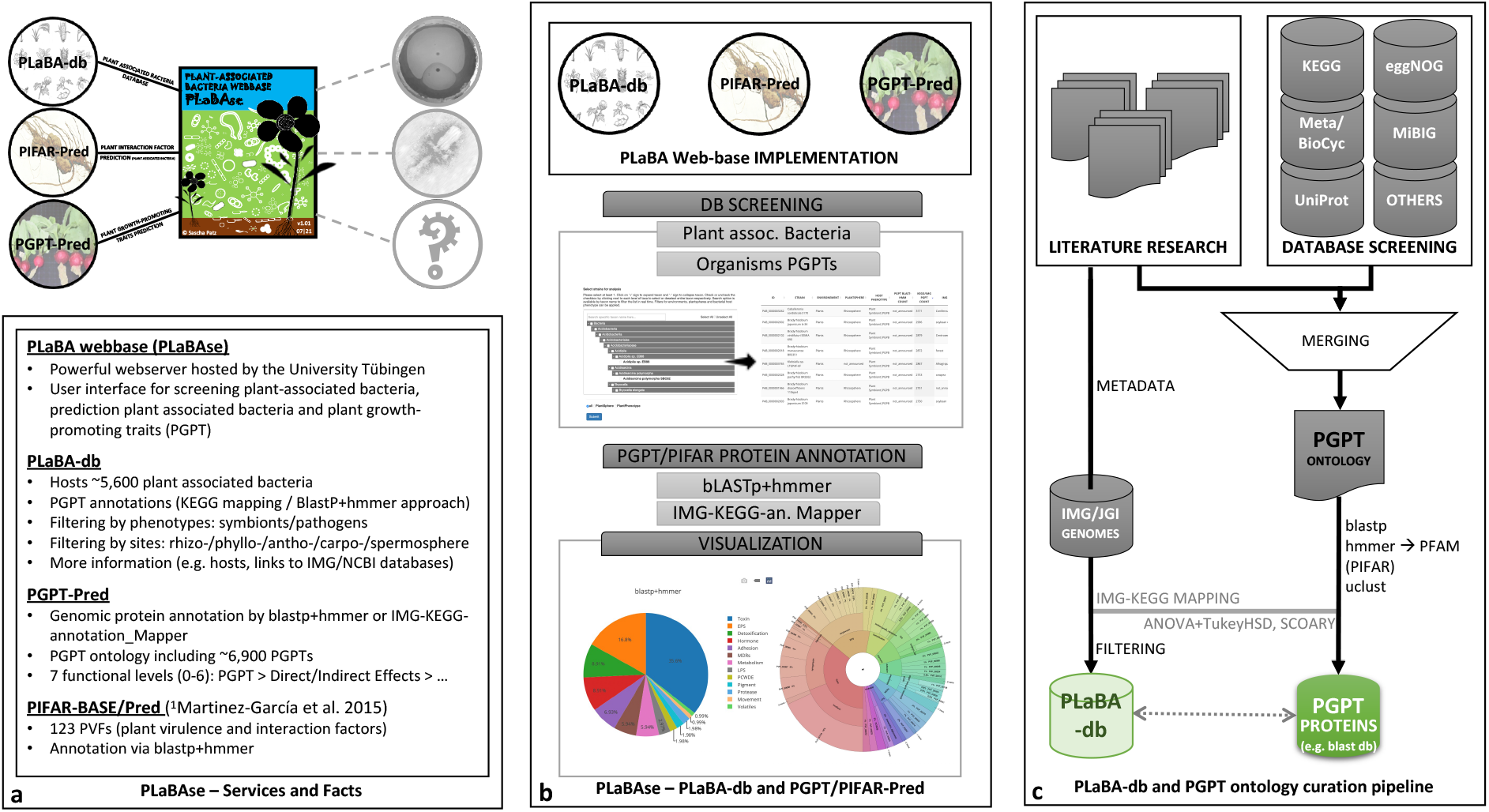
PLaBAse curation workflow and implemented features. (c) Summary of PLaBAse platform with short descriptions of the services (databases and tools). (b) Preview of the various strategies for strain and functional screening of known or novel bacterial genomes and their result data visualization. The blastp+hmmer approach allows direct annotation of protein FASTA files, whereas the IMG-KEGG-annotation Mapper accepts IMG-KEGG annotation files as input. The data can be visualized as pie chart and krona plot. (c) Overview of the independent approaches to establish (i) the PLaBA-db, by fetching genomes and their metadata from the IMG-Server and filtering for plant-associated bacteria only, and (ii) the PGPT ontology, curated from literature-derived PGP processes and protein families. Respective PFAM domains are attached to the PGPT ontology and proteins are stored in a blast database.

Our web resources are compatible with Mozilla Firefox (v90.0.1), Google Chrome (v91.0.4472.164), Safari (v14.1.1) and Edge (v91.0.864.67).

### Ontology Construction for PGPTs

The definition of bacterial PGPTs (protein families) and their hierarchical functional classes is based on publicly available literature, including Glick et al. 1996 and 2012 ^56,68^, De Souza et al 2015 ^16^, Gouda et al 2018 ^57^, Suarez et al 2019 ^69^, Afzal et al 2019 ^58^ and Compant et al 2021 ^70^, and various OMICs databases (see Data Acquisition and Storage of PGPTs). As PGPT in our ontology we considered each genomic element (gene encoded protein), that was described as fulfilling a PGP function in the literature and that could be assigned to at least one PGPB (see Introduction, Suppl. Table 1). In a few cases we added manually curated PGPTs only if they could be assigned to a major PGPT functional pathway and thus might be present in an as of yet unknown symbiotic bacteria or acquired by horizontal gene transfer. Approximately 6,900 curated PGPTs were organized into a hierarchical (tree-like) structure that represents each trait (protein family) as a single PGPT at the leaf nodes, assigned with a PGPT classifier. Each classifier represents a unique PGPT identifier (e.g., PGPT0000005), one gene symbol or gene name (e.g., *nif*B) and the corresponding functional KEGG, COG and PFAM annotations. Internal nodes indicate respective PGPT functional classes towards the root node, termed “PGPT”. The tree has seven functional levels (level 0 to 6) and is stored in Newick format as displayed in Figure 1a,b and Figure 2. Because PGPTs can be assigned to various plant-beneficial classes on internal levels, the tree is considered a multi-labeled tree at the level of leaves.

**Figure 2:**
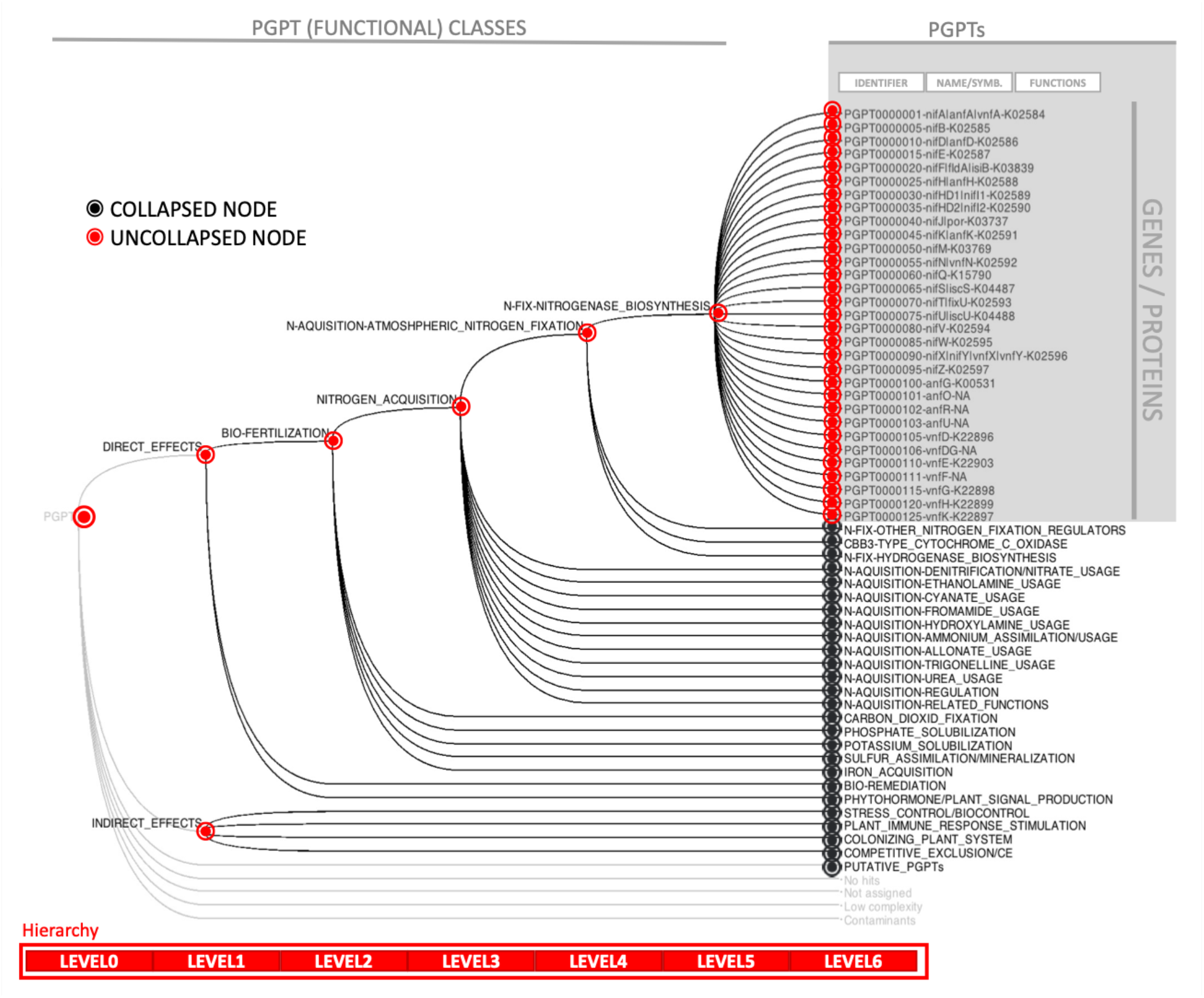
PGPT ontology as hierarchical multi-labeled tree. PGPT-BASE comprises currently 6,912 unique PGPTs. These are classified on six functional subclass levels (level 0-5). For example, the PGPTs encoding the nitrogenase complex (level 5), needed for atmospheric nitrogen fixation (level 4), are shown up to leaf node level 6, assigned to their sub-classes towards the root node (red dotted path of uncollapsed nodes). The tree visualization was produced using MEGAN6.

### Data Acquisition and Storage of PGPTs

PGPT-related bacterial protein sequences for all classifiers were obtained from various functional annotation systems (Figure 1 c), namely KEGG (Kyoto Encyclopedia of Genes and Genomes, https://www.kegg.jp), GO (Gene Ontology Resource, http://geneontology.org), eggNOG (http://eggnog5.embl.de/#/app/home), UniProt (https://www.uniprot.org) and Meta-/Bio-Cyc (https://biocyc.org, https://metacyc.org). Information about secondary metabolites that have direct or indirect PGP relevance, were manually added by screening MIBiG compounds (https://mibig.secondarymetabolites.org). Critical cases, where appropriate sequences could not be identified, also not by genomic location, taxonomic assignment, and associated pathways, were discarded in the current version of PGPT classification. By clustering all identified PGPT protein sequences with the tool uclust (v1.2.22q), redundant sequences per PGPT classifier were removed to obtain a set of non-redundant PGPT reference sequences:

~~~
uclust --sort <PGPTid-proteins.faa> --output <PGPTid-prot_sor.faa>
uclust --input <PGPTid-prot_sor.faa> --uc <PGPTid-proteins_res.uc> --id 1.00 –amino
uclust --uc2fasta <PGPTid-proteins_res.uc> --input <PGPTid-prot_sor.faa> --output
<PGPTid-proteins_non-red.faa> --types S
~~~

The remaining ∼6,965,955 protein sequences were indexed in a blastp database and carry their PGPT identifier in the sequence header.

~~~
makeblastdb -in <PGPTid-proteins_non-red.faa> -title pgpt_db_Aug2021 -parse_seqids
-dbtype prot
~~~

Additionally, PFAM domains were identified using hmmsearch (http://hmmer.org/, HMMER 3.3) and together, with PGPT protein identifiers, saved in the PGPT ontology:

~~~
hmmsearch --tblout <PGPTid-prot_unique.pfam> -E 1e-5 Pfam-A.hmm
PGPTid-proteins_non-red.faa
~~~

The Pfam-A.hmm file was retrieved on 2021/08/13 via ftp server:

~~~
wget ftp://ftp.ebi.ac.uk/pub/databases/Pfam/releases/Pfam34.0/Pfam-A.hmm.gz
~~~

### Genomic Data Acquisition and Classification

Genomic and metadata information of 71,931 bacteria listed in the IMG database (https://img.jgi.doe.gov), were downloaded in November 2020 ^71,72^. Sequences were grouped (i) by affiliation to taxonomic levels (phyla, class, order, family, genus, species), (ii) by isolation sites (plant, human, animal, soil, aquatic, food, industrial, not_defined) and (iii) by either pathogenic or symbiotic phenotype (Table 1). The industrial group comprises strains isolated from sewage plants, slurry, and bioreactors. The obtained data were extended by information listed in the NCBI RefSeq assembly database (https://ftp.ncbi.nlm.nih.gov/genomes/refseq/bacteria/assembly_summary.txt), and on the KEGG repository (https://www.genome.jp/kegg-bin/download_htext?htext=br08601_key.keg). For downstream analysis of PGPTs, only the functional annotations of those 70,540 genomes that have at least 450 genes, were further considered. The limit was estimated using the current known minimal gene count of *Mycoplasma mycoides* that comprises 473 genes ^73^. Finally, 38,735 genomes could be assigned to one of the metadata groups mentioned above and were processed further. All annotated bacterial genomes were grouped according to their environmental isolation sites, their plant spheres (organs) and known bacteria-induced plant symbiotic or pathogenic phenotypes (Table 1, Suppl. Tables 2-4). Thus, in total 5,565 plant-(PA), 17,046 human-(HA), 5,065 animal-(AA), 5,464 soil-(SA), 5,656 aquatic-(QA), 1,190 food-(FA) and 1,166 industrial-associated (IA) strains were processed, and the genus diversity calculated (Table 1 left-hand). Among them, the datasets for QA, SA and PA had highest genus diversity, albeit approximately three times higher strain counts were benchmarked for HA. Due to our explicit interest in plant-associated bacteria, we determined 2,261 rhizospheric (PA_RHIZ) and 708 phyllospheric (PA_PHYL) bacteria (Table 1 right-hand, upper part), beside other strains related to under-represented plant environments, like the anthosphere (PA_FLOW: 31), carposphere (PA_FRUIT: 55) and spermosphere (PA_SEED: 134). For the bacteria-induced host phenotype, there are 855 plant symbiotic and 890 phytopathogenic strains (Table 1 right-hand, bottom part).

**Table 1.** Overview of the classified bacterial strains on JGI/IMG used for analysis of PGPBs/PGPTs^1^.

| Isolation site | Unique strain count | Genus diversity (Simpson Reciprocal I.) | Plant Spheres \|\| Phenotypes | Unique strain count |
| --- | --- | --- | --- | --- |
| Plant-associated (PA) | 5,565 | 233.84 | Rhizosphere (PA_RHIZ) | 2,261 |
| Human associated (HA) | 17,046 | 26.11 | Phyllosphere (PA_PHYL) | 708 |
| Animal associated (AA) | 5,065 | 70.23 | Anthosphere (PA_FLOW) | 31 |
| Soil associated (SA) | 5,464 | 608.67 | Carposphere (PA_FRUIT) | 53 |
| Aquatic associated (QA) | 5,656 | 948.81 | Spermosphere (PA_SEED) | 134 |
| Food associated (FA) | 1,190 | 23.89 | Plant symbionts (PA_SYMB)<sup>2</sup> | 855 |
| Industrial associated (IA) | 1,166 | 11.05 | Plant pathogens (PA_PATH) | 890 |

<sup>1</sup>source: https://img.jgi.doe.gov, total unique strain/genome count with informative metadata: 38,735
<sup>2</sup>among them only 248 labelled as “PGPB” by publication title; source: https://www.ncbi.nlm.nih.gov/genbank/, https://pubmed.ncbi.nlm.nih.gov

A One-Way-ANOVA with TukeyHSD test was performed to calculate statistically significant differences (p-value < 0.05) for genome size and gene count between all groups (R packages: dplyr, ggpubr).

### Bacterial Genome-Wide PGPT Enrichment Study

As a sanity check to expose any annotation biases, we first mapped each IMG-genome functional KEGG identifier set against the PGPT ontology, applying the IMG-KEGG-annotation_MAPPER via the PGPT-Pred tool on our website. The respective PGPT annotation for currently 5,565 plant-associated bacteria including the strains meta-information (e.g., plant sphere) are accessible via PLaBA-db.

In a next step genome-wide enrichment statistical analysis was performed using Scoary v.1.6.16 (https://github.com/AdmiralenOla/Scoary) to examine the enrichment of single PGPT proteins up to complete PGPT functional classes ^74^. Scoary was executed with the recommended command for such an analysis:

~~~
scoary -g <PGPT file> -t <metadata file> -p 0.05 -c BH -w --no_pairwise
~~~

We analyzed (i) the isolation site-specific enrichment of PGPTs and subsequent (ii) the PGPT functions, as well as their enrichments in plant-associated and symbiotic bacteria (including PGPBs). Based on Fisher’s exact tests and Benjamini-Hochberg corrections, the odds ratio was calculated and reported only if p-values were smaller than a 0.05 significance threshold. Python modules matplotlib, numpy and pandas and the R library “gplots” were used for plotting all results as boxplots and/or clustered heatmaps (heatmap.2 function, Pearson correlation distance with average linkage). As a negative control for PGPTs, we added the human pathogenicity island (*cag* genes), which was expected to be absent in plants but present in human-associated strains. It was discarded from the PGPT ontology after analysis as the entire gene cluster could not be found in plant-associated strains.

## RESULTS

In the past decades scientists have isolated bacterial strains with high plant-beneficial effects, which harbor a wide range of PGP functions. Unfortunately, explicit indication of such strains is missing in most databases. While terms like “pathogen” are extensively used, PGP-related statements, like “plant endosymbiont” or “bioremediation” occur much less frequently. Regarding the literature, plant growth-promoting bacteria might be abundant in various phyla, but often only one or a few strains per genus are described as PGPB. This hampers genome-wide association studies in a pan-genome manner aimed at determining universal traits and genes linked to plant growth-promotion. Varying PGP potential among strains due to different sets of PGP traits (PGPTs; genes/proteins) currently hinders consistent identification of PGPBs. Intensive literature research was needed to collect all current information about PGP strains and PGPTs with their functional classes. However, differentiation between plant symbionts and PGPBs on a phenotype and genotype level remains a challenge and is perhaps not feasible, so that we consider here both as plant-associated symbiotic bacteria (PA_SYMB) together. Thus, we do not yet provide a PGPB prediction tool, but an important step towards it, by providing a comprehensive web resource for analysis of plant-associated bacteria (PLaBAse), aimed at ensuring standardized annotation of main PGP processes on a genomic and proteomic scale.

### The PLaBAse-Web resource and services

The web resource for analysis of plant-associated bacteria, short PLaBAse (Figure 1a,b), is a platform that provides (i) a MySQL database of 5,565 plant-associated bacteria (PLaBA-db), selected from 70,540 genomes, that were analyzed in this work, (ii) a prediction tool for bacterial plant growth-promoting traits (PGPT-Pred), and (iii) a tool for prediction of plant-associated bacteria (PIFAR-Pred). For all plant-associated bacterial strains the PLaBA-db stores PGPT annotations, isolation sites (environments, plant spheres, bacteria-induced host phenotypes), taxonomic lineage, genomic features, and respective links to the IMG and NCBI databases. Taxon- and isolation site-specific filters allow selection of specific bacterial strains. Both tools, PGPT-Pred and PIFAR-Pred, perform genomic annotation of protein sequences of single genomes. One can submit protein FASTA sequences by copy-paste or upload. The input data undergoes an initial sanity check, and then a job is queued, and the user is notified via email about submission and completion status. Alternatively, the IMG-KEGG-annotation_Mapper feature of the PGPT-Pred tool enables upload of an IMG-derived KEGG annotation file, or its content, and assigns proteins with K numbers to the PGPT ontology. After completion of the job the user is provided with a link to a result page via email, which is available for 15 days. Results are provided to the user in form of interactive plots, e.g., as pie charts or KRONA plots, and as tab-delimited text files. Here, the user can choose between blastp+hmmer hits or blast-only hits, as novel isolated strains might harbor genes that were not considered when building up current PFAM-domain HMM-models.

More detailed information about the PGPT ontology implemented in the PGPT-Pred tool is shown in Figure 1c. Its capability to provide standardized PGPT annotation of bacterial genomes across various isolation sites will be presented in the subsequent results.

### The PGPT Ontology System

The PGPT ontology is a literature-derived classification of PGP functions (Figure 1c), represented by a functional tree (Figure 2, Suppl. Table 1), where each single PGPT, i.e., a protein family, is represented as a leaf node. Their assignment to more general functional classes is achieved by moving up the internal nodes toward the root node. Because individual traits may fulfill multiple functions, a single trait may occur more than once as a leaf node, so that the tree must be regarded as multi-labeled. The 6,912 PGPTs currently listed in our ontology are assigned to 12,593 leaf nodes (level 6). At a leaf node each PGPT classifier consists of a PGPT identifier, the gene name or symbol, and an information vector, storing KEGG, eggNOG and PFAM annotations. There are six functional sublevels, comprising the root node (“PGPT”), three PGPT classes on level 1, eight PGPT classes on level 2, 47 classes on level 3, 280 classes on level 4 and 1,966 PGP classes on level 5 (Table 2). We anticipate that these numbers will change in future releases. On level 1 we distinguish mainly between (i) direct or (ii) indirect PGP effects, whereas o level 2 major PGPT classes, like bio-fertilization, bioremediation, phytohormone/plant signal production, stress control (e.g., biocontrol), plant immune response stimulation, colonizing plant system and competitive exclusion are listed. In total, 6,965,955 protein sequences are assigned to all the PGPTs, currently, and are not restricted to just plant-associated bacteria, to estimate the PGPT occurrences for strains of other environments as well.

**Table 2.**
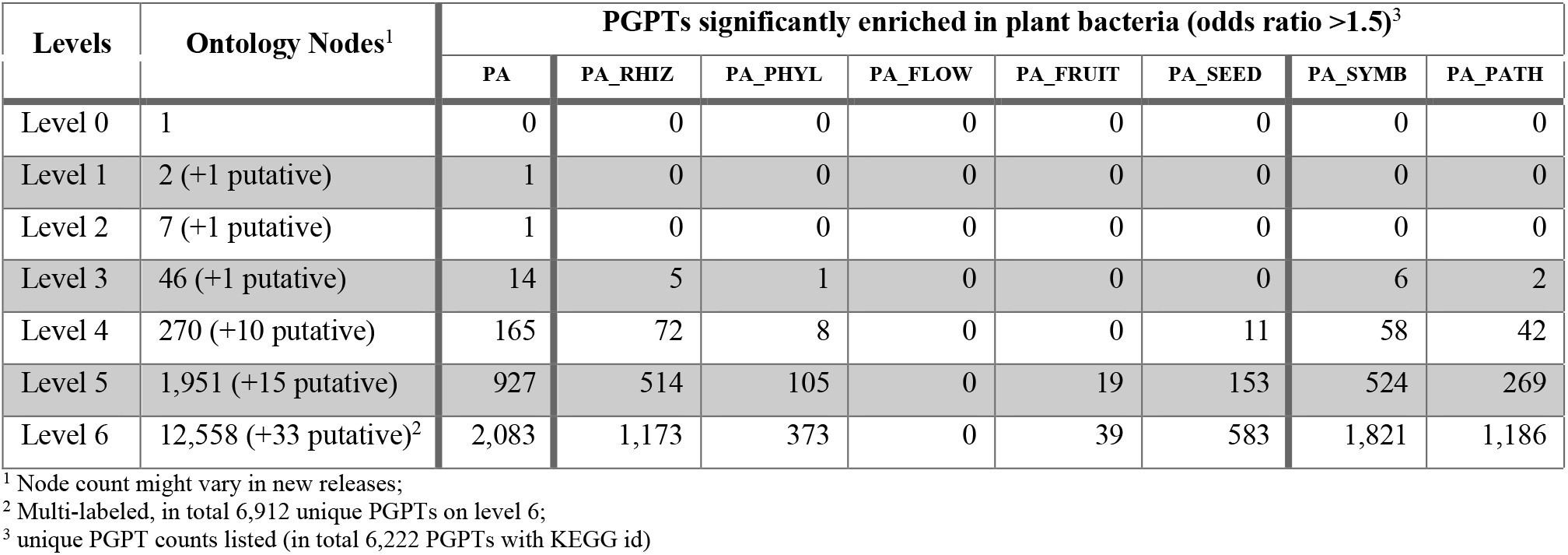
Summary of curated PGP functions represented in the PGPT ontology, and their enrichment based on KEGG-PGPT mapping.

| Levels | Ontology Nodes<sup>1</sup> | PGPTs significantly enriched in plant bacteria (odds ratio >1.5)<sup>3</sup> |  |  |  |  |  |  |  |
| --- | --- | --- | --- | --- | --- | --- | --- | --- | --- |
|  |  | PA | PA_RHIZ | PA_PHYL | PA_FLOW | PA_FRUIT | PA_SEED | PA_SYMB | PA_PATH |
| Level 0 | 1 | 0 | 0 | 0 | 0 | 0 | 0 | 0 | 0 |
| Level 1 | 2 (+1 putative) | 1 | 0 | 0 | 0 | 0 | 0 | 0 | 0 |
| Level 2 | 7 (+1 putative) | 1 | 0 | 0 | 0 | 0 | 0 | 0 | 0 |
| Level 3 | 46 (+1 putative) | 14 | 5 | 1 | 0 | 0 | 0 | 6 | 2 |
| Level 4 | 270 (+10 putative) | 165 | 72 | 8 | 0 | 0 | 11 | 58 | 42 |
| Level 5 | 1,951 (+15 putative) | 927 | 514 | 105 | 0 | 19 | 153 | 524 | 269 |
| Level 6 | 12,558 (+33 putative)<sup>2</sup> | 2,083 | 1,173 | 373 | 0 | 39 | 583 | 1,821 | 1,186 |

<sup>1</sup> Node count might vary in new releases;
<sup>2</sup> Multi-labeled, in total 6,912 unique PGPTs on level 6;
<sup>3</sup> unique PGPT counts listed (in total 6,222 PGPTs with KEGG id)

### Plant- and soil associated bacteria have higher genome sizes across various environments

We observed a significantly higher genome size and gene count for strains associated to PA and SA with an average of 5,312 – 5,344 genes distributed across 5,576,974 – 5,665,934 nt, compared to HA (3,869 genes; 3,936,973 nt), AA (3,201 genes; 3,291,610 nt), QA (4,113 genes; 4,325,167 nt), FA (3,482 genes; 3,507,913 nt) and IA (4,492 genes; 4,772,482 nt) (Figure 3a,d; Suppl. Table 2). The largest genomes were measured for plant- and/or soil-associated bacteria, such as *Mumia* flava MUSC 201, *Streptomyces* sp. S1D4-11, and *Saccharothrix* sp. ALI-22-I. A possible bias due to unequal distribution of incomplete genomes across groups can be excluded, as similar and even stronger patterns were observed when considering complete genomes only (Suppl. Figure 1).

**Figure 3:**
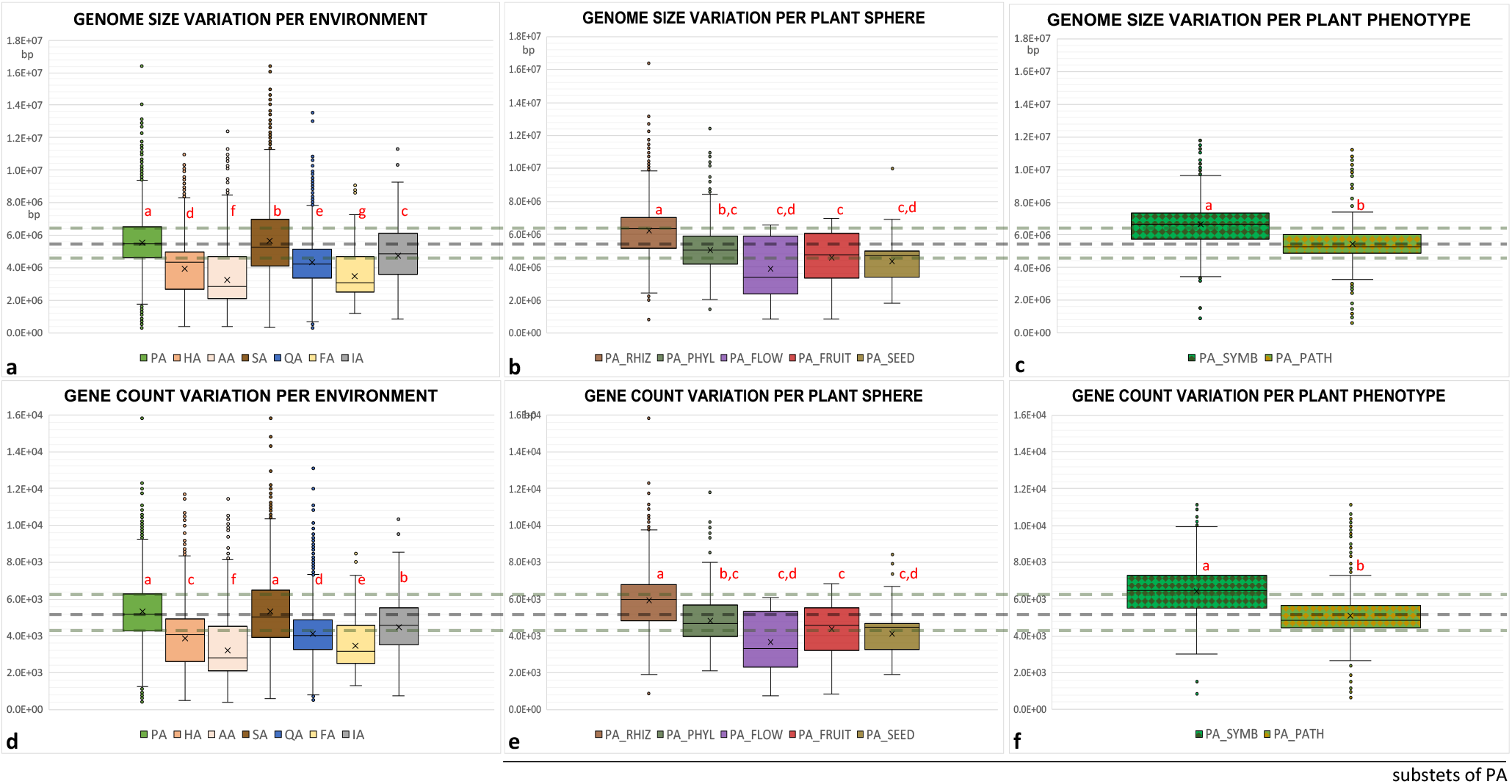
Genomic features of bacterial strains in association to isolation site and plant-specific classes stored in the JGI/IMG database. Upper plots show variations in genome sizes. Bottom plots display gene-count variation dependent on isolation site (a,d), plant sphere (b,e) and bacteria-induced plant phenotype (c,f). Dotted lines highlight the median and percentiles of plant-associated bacteria. The sample count per group is listed in Table 1. Significant differences with an adjusted p-value < 0.05 to each other per plot were tested by a One-Way-ANOVA with TukeyHSD and indicated by red letters in each plot. (PA plant-associated, HA human ass., AA animal ass., SA soil ass., QA aquatic ass., FA food ass., IA industrial ass., PA_RHIZ plant rhizosphere, PA_PHY plant phyllosphere, PA_FLOW plant anthosphere, PA_FRUIT plant carposphere, PA_SEED plant spermosphere, PA_SYMB plant symbiotic bacteria, PA_PATH pathogens.)

### Across different plant spheres the higher genome sizes are related to rhizospheric bacteria

In the context of plant sphere-specific strains, as subsets of PA, we observed higher values for both, genome size and gene count, in rhizospheric strains (5,913 genes; 6,198,683 nt), that exceed the average of all PA strains together, and compared to phyllospheric (4,840 genes; 5,057,835 nt), anthospheric (3,678 genes; 3,895,520 nt), carpospheric (4,386 genes; 4,594,780 nt) and spermospheric (4,143 genes; 4,362,472 nt) isolates (Figure 3b,e; Suppl. Table 3). Anthospheric bacteria, show the lowest gene count on average, but comprise a large number have a count on par with the superordinate PA average. Seed isolates comprise a large proportion of strains with counts around the PA 25% percentile, below PA average.

### Overall highest genome sizes are found in plant-symbiotic bacteria

Regarding the bacteria-induced plant phenotype, caused by symbiosis or pathogenicity, we observed highly significant differences for respective bacterial groups (subsets of PA). Plant symbiotic bacteria have larger genomes that encode more genes on average (6,644,481 bp and 6,443 genes) compared to pathogenic strains (5,447,664 bp and 5,090 genes) and compared to the superordinate PA (Figure 3c,f; Suppl. Table 4). As a side note, human and animal pathogens together (2,609 strains) have on average a much lower genome size of 3,305,759 bp and fewer genes counts of 3,255 (not shown). Similar results have been obtained for coding sequences (CDS) only (not shown). Strain specific variabilities and outliers were detected for all groups. Summarizing the results in brief, symbiotic plant-associated bacteria harbor more genes and have larger genomes. This is mainly due to rhizospheric bacteria and is apparent for SA bacteria, too.

### Environments: Higher PGPT counts mostly associated with plant- and soil associated bacteria (Level 0 and 1)

To illustrate the utility of the PGPT ontology, we analyzed the PGPT distribution across multiple bacterial genomes in a standardized manner and evince the contribution of PGPTs on the observed genome size and gene count variations. Figure 4a depicts the PGPT abundance per functional level, compared in the order from left to right: (i) all seven environmental isolation sites, (ii) five plant spheres, and (iii) between plant symbionts and plant pathogens. Using the genomic IMG-KEGG annotation mapping against the PGPT database, comprising 6,222 PGPTs with a K number, we detect overall an on average significant higher PGPT count per genome on level 0 for the environmental isolation sites PA (∼1,749 PGPTs) and SA bacteria (∼1,559) compared to all other environments. However, we also observe counts around the PA average for several strains designated as HA, FA and IA, although their average counts are mostly lower than the 25% quantile for PA (Figure 4a, upper left plot). On PGPT functional level 1, where we distinguish between the three classes: direct or indirect effects, and putative PGPTs (Figure 4a, plots of the three bottom rows), similar patterns, as discussed for level 0, occur across all sites. However, the highest overlap between PA and in particular HA strains are seen predominantly by indirect traits. Interestingly, when considering the gene count per genome, we can estimate a stabile PGPT ratio of approximately one third for all environments.

**Figure 4:**
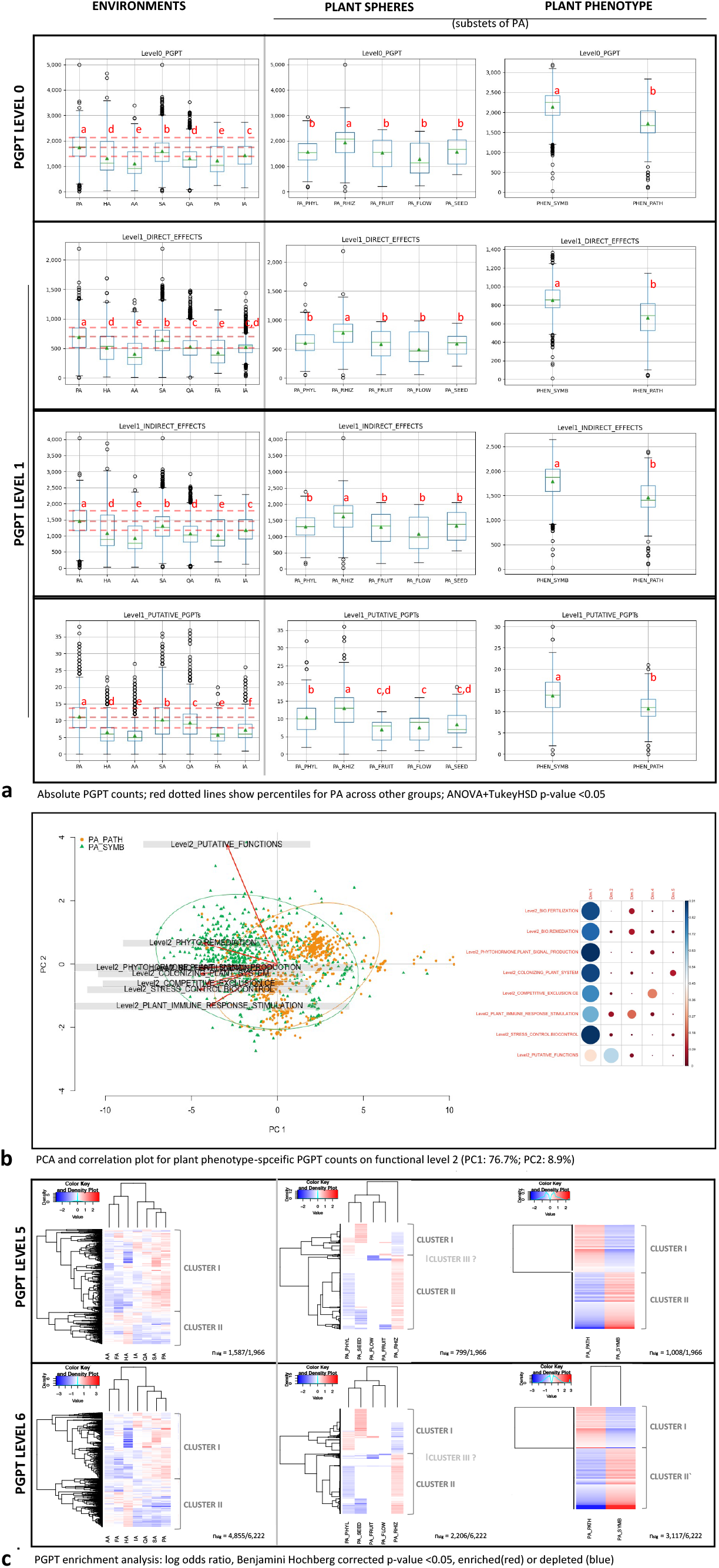
Environmental, plant sphere and phenotype specific distribution of PGPTs. (a) Boxplots of absolute PGPT counts across seven bacterial environments, five plant spheres and two bacteria-induced plant phenotypes (abbreviation see Figure 3) for functional level 0 and 1 of the PGPT ontology (red dotted support lines allow better comparability of PA quantiles). Statistical significances were established with ANOVA and TukeyHSD (p-value <0.05) specified by red letters. (b) PCA biplot (Bray-Curtis clustering) and correlation matrix for plant symbiont (green) and pathogens (orange) specific PGPT counts on level 2. (c) PGPT enrichment analysis on level 5 and level 6 (sole PGPT protein families) on y-axis (for terms see Suppl. Tables 6 and 7) by clustering of respective PGPTs count log odds ratios. The scale varies between the plots but specifies depletions in blue and enrichments in red color gradations with the darkest colors as boundaries.

### Plant spheres: Highest PGPT counts are strongly related to rhizospheric bacteria (Level 0 and 1)

Within different plant spheres (Figure 4a, upper centric plot), the highest average count of PGPTs per genome on level 0 is found for rhizospheric strains (∼1,943), while the lowest average occurs for the anthospheric strains (∼1,314). But again, in total we observe an overlapping section of PA strains with higher PGPT counts among all plant sites. On level 1 we see similar patterns for direct and indirect effects.

### Bacteria-induced plant phenotypes: On average the highest PGPT counts for plant-symbiotic bacteria (Level 0 and 1)

More importantly, an on average significantly higher count of ∼2,139 PGPTs per genome (level 0) is measured for plant symbiotic strains compared to ∼1,725 PGPTs per genome for pathogens (Figure 4a, upper right plot). Noteworthy, when considering all pathogenic strains, including human and animal pathogens, the value is decreases to ∼1,300 PGPTs per genome (not shown). Similarly, on level 1 symbionts reveal the highest average PGPT counts for each class. The highest overlaps between both phenotypes (symbiotic and pathogenic) were observed for indirect effects.

### All level 2 PGPT functions show on average higher counts for plant-associated, plant rhizospheric and plant symbiotic strains

Similar patterns show up for all eight subclasses on level 2, namely bio-fertilization, bioremediation, phytohormone and plant signal production, stress control / biocontrol, colonizing plant system, competitive exclusion, plant immune response stimulation and putative PGPT functions (Suppl. Figure 2); the highest average counts are assigned to PA (followed by SA), rhizobacteria and plant symbionts. For plant symbionts, the highest difference on average to pathogens was detected for colonization with a count for symbiotic strains of ∼1,170 PGPTs per genome. We observed high counts for bio-fertilization, stress control / biocontrol, colonization, and competitive exclusion in several strains of other environments as well, albeit the group averages were quite low, especially in human-associated strains.

Thus, in brief, overall high counts of PGPTs on level 0 to 2 are strongly associated with, but not restricted to PA and known plant symbionts.

### PGPTs facilitate differentiation between Plant Symbionts and Pathogens on Level 2

Altogether, we demonstrate in Figure 4b that the novel ontology covers the main genetic drivers for plant symbiotic strains, including PGPBs (Suppl. Figure 3). 77% of the variance in the PGPT count between symbionts and pathogens can be explained by higher PGPT assignments to all PGP classes on Level 3 (Suppl. Figure 4a, b). In addition, 9% of the variance between pathogenic and symbiotic strains can also be explained by mostly putative functions, where the highest variance is associated with the known PGPB *Nitrospirillum amazonense* CBAmc, among others. However, while the majority can be distinguished on the level 2 class PGPT count, single symbiotic and pathogenic strains appear in both clusters.

### PGPT Enrichment Analysis for level 3 to 6

Due to the previous observation and to the increasing, and thus here not fully presentable, amount of PGPT classes from level 3 to level 6, we performed a genome-wide enrichment study to figure out which PGPT functional subclasses (level 3-5) and single PGPTs (level 6) are significantly enriched in PA, in different plant spheres and, especially, in plant symbionts (Suppl. Tables 5-7). As the results for all mentioned levels are quite similar, only level 5 and 6 are displayed as clustered heatmaps in Figure 4b. Both levels allow a very detailed view into functional PGPT groups and single PGPT genes, respectively.

### Environmental specific clustering of PGPTs (Level 5 and 6)

When focusing on environments, two distinct clusters confirm the previously observed PGPT count overlap also on enrichment for level 5 classes and sole PGPTs on level 6. Cluster I represents PGPT enrichments for the environmental isolation sites PA, SA, QA (and slightly IA) that appear to be depleted in HA, AA and FA. The pattern is vice versa for cluster II (Figure 4c left plots). Thus, in total 2,188 of 4,855 significantly altered PGPTs on level 6 are enriched in PA (cluster I), with an odds ratio ≥1.5 (Table 2). Interestingly, 1,357 PGPTs of these are also enriched in SA bacteria, that reveal in total 2,019 enriched PGPTs. However, 866 PGPTs show significantly higher enrichments in PA only (Suppl. Table 5). Up to 74 traits (including the negative control: *cag* genes) were absent in plant-associated strains analyzed in the given dataset, associated mostly to indirect effects like competitive exclusion and colonization. Among the remaining potential PGPTs that appear to be less abundant, but present, in PA bacteria (cluster II, level 6), we found nitrogenase-associated *nif*SIJU and *nif*HD1/2 genes and hydrogenase genes *hyf*A-R, *hyp*A-F and *hup*SL/*hup*C, that are known to fulfill also other functions beside their incorporation in the atmospheric nitrogen fixating *nif* gene cluster and / or co-occurrence in some PGPBs.

### Plant-sphere specific clustering of PGPTs (Level 5 and 6)

Considering plant spheres, more than 2/3 of all enriched classes and single PGPTs (level 5 and 6) are related to rhizospheric strains (1,173 of 2,206 significantly altered PGPTs; cluster II) and are depleted or not significantly different in other spheres (Figure 4c centric plots, Table 2, Suppl. Table 6). Although we must consider a bias due to higher strain counts of rhizospheric bacteria, we see distinct PGPT enrichments for the less abundant spermospheric (583), phyllospheric (373) and slightly carpospheric (39) isolates (cluster I) only. In particular, seed isolates are enriched in PGPTs related to plant-derived compound transport, PTS systems, pilus systems, surface attachment genes, peptidoglycan and lipoprotein remodeling, quorum sensing and biofilm regulation, amylovoran biosynthesis, osmotic and nitrosative stress tolerance, biosynthesis of vitamin B1, B2, B5, B6, B9, C, K, GABA, carotenoids and acetoin / butanediol volatiles, iron transport systems and siderophores, heavy metal and xenobiotic resistance, and nitrogen uptake and sensing. Interestingly, phyllosphere specific gene clusters are related to nitrate reduction (*nfr*ABCDEG), siderophore and iron transport (e.g., *irp*1-5, *bae*SR), pilus system (e.g., *pil*B-Z), type III and IV secretion systems (e.g., *hpa*ABC, *hpa*1-2, *dot*BCD, *icm*BEGKLOT), phyto hormones and plant signaling compounds (e.g., *crr*CDEFIW, *isp*ACDF, *men*ABCDEF). A third possible cluster was detected, comprising mostly depleted traits, which is specific to anthospheric strains. These 38 significantly depleted traits comprise genes related to organic acid pathways, potassium uptake, (siro-) heme and molybdenum biosynthesis.

### Plant-symbiotic and -pathogenic bacteria reveal distinct PGPTs enrichment (Level 5 and 6)

Regarding bacteria-induced plant phenotypes, two distinct cluster were found with higher amount of PGPTs enriched for symbiotic (cluster II), rather than pathogenic bacteria (cluster I) on level 5 and 6 (Figure 4c right plots, Table 2, Suppl. Table 7). In total 1,821 level 6 PGPTs are significantly enriched for symbiotic strains (odds ratio >1.5) and are not shared with the enriched ones of pathogens (1,186). Interestingly, 393 of the enriched symbiosis-associated PGPTs were not found to be enriched in PA, when PA was compared with other environments on level 6. These enriched plant symbiotic traits can be associated with colonization (30%), competitive exclusion (21%), bio-fertilization and stress control / biocontrol (13-15%), bioremediation (11%), phytohormone / plant signal production (8%), and plant immune response stimulation or putative traits (0.4-1%). High counts of symbiotic traits were found for plant-derived substrate usage (e.g., degradation and transport of C4-dicarboxyrate, cellobiose, erythritol, glucose, fructan, fructose/fructose-lysine, mannitol, nicotinate, raffinose, rhamnose, stachyose, starch, tagatose, tricarboxylate), nitrogen acquisition (e.g., genes encoding for atmospheric nitrogen fixation, nitrate reduction, glutamate metabolism and transport), CO2 fixation by RuBisCo, plant vitamin production (e.g., vitamin B1, B2, B5, B6, B9, B12, K), plant hormone metabolism (e.g., abscisic acid degradation, cytokinin, xanthine / carotenoid / terpenoid metabolism, GABA and phospholipid biosynthesis, spermidine / putrescine metabolism, plant signaling volatiles), neutralizing abiotic stress (e.g., nitrosative / oxidative, osmotic or salinity stress) xenobiotic and heavy metal remediation, phosphate solubilization (e.g., phosphonate degradation or degradation by sulfuric acids or diverse organics acids), neutralization of biotic stress (e.g., bactericidal or fungicidal compounds), colonization-surface attachment (via e.g., lipids, adhesins, teichoic/colonic acids, vitamin B7), sporulation, transposases or genes related to circadian rhythm or photosynthesis electron transfer (Suppl. Figure 4). In contrast, in phytopathogens enriched PGPTs are type III secretion system secreted proteins (encoded by genes *hrp*E, *hpa*1, *hpa*2, *hpa*AB), type Vb autotransporter secretion (*big*A), quorum sensing PAME regulated exopolysaccharide production (*eps*EF), quorum sensing DSF precipitation and signaling (*rpf*G), host invasion factors (*sip*AC/*ipa*AC), surface attachment lipid and lipopolysaccharide biosynthesis and modification (*lpx*R, *ops*X, *waa*RTY/*rfa*Y) and yersiniabactin siderophore biosynthesis (*irp*1-5), among others. The flagellar assembly encoding gene clusters are both enriched in pathogenic strains, but while the *flg* gene cluster is slightly enriched, the *fli* gene cluster shows a higher odds ratio. Excitingly, the flagellin encoding gene (*fli*C), the main flagellum component and immunity response trigger, is slightly enriched in phytopathogens, but is more enriched in symbionts when comparing to the huge amount of human and animal pathogens (not shown), as well. The same pattern is present for phosphate solubilization by gluconic acid production and secretion (*pqq* genes).

## DISCUSSION

A systematic analysis of the distribution of plant growth-promoting traits (PGPTs) among the bacterial kingdom, considering strains environmental, plant sphere, and bacteria-induced plant phenotypes (symbiotic or pathogenic) properties, will improve the identification and case- or site-directed application of plant growth-promoting bacteria (PGPBs) as beneficial bio-inoculants in plant production systems. Their beneficial potential in bio-fertilization, plant growth regulation, bioremediation, biotic and abiotic stress control (e.g., biocontrol) has been investigated in previous research of various scientists and contributed to the achievements of that study, as stated below ^23,75–77^.

Our PLaBAse web resource aims to facilitate (i) the search of known plant-associated bacterial strains, especially plant symbionts and PGPBs, and their PGPTs using PLaBA-db, (ii) the prediction of plant-associated bacteria using PIFAR-Pred, and (iii) standardized annotation of PGPTs in bacterial genomes using PGPT-Pred. The latter is based on the PGPT ontology, which organizes most known plant-beneficial bacterial traits (genes or proteins), according to various plant-related functional classes in a hierarchical style, a characteristic of other most functional databases and ontologies (such as VFDB or GO ^28,78^) ^16,56–58,68–70^. All PGPTs can be analyzed on seven different levels (level 0-6), providing different levels of functional resolution.

Currently, plant-beneficial traits related to Archaea are not stored in our ontology, because such traits are rarely described in literature, perhaps due to the fact that Archaea appear to be less abundant in and on plant tissues, or due to their isolation biases ^79^. We are also aware that certain classes, such as for host colonization or competitive exclusion, might be essential for interaction with other hosts or environments rather than only with plants ^80^. Indeed, the application of the PGPT ontology to bacterial genomes of various environments helps understand the distribution of PGPTs.

In general, PGPTs in PA genomes can be grouped into three patterns, namely (i) unique, (ii) enriched, and (iii) those present, but not enriched. The latter pattern confirms that some PGPTs are not restricted to plant-associated bacteria and do not only contribute to PGP. Even more obvious, among the PGPTs enriched in plant symbionts, 393 were not enriched in PA genomes, when compared to other environments, suggesting their varied roles, but especially in plant growth-promotion.

The distinct clustering detected for significantly enriched or depleted PGPTs in plant symbionts on most levels might also point to their intrinsic agent and implies a different weighting of PGPTs and associated plant-beneficial functions. The dominance and diversity of genes that are important for colonizing the plant and for competitive behavior might give one explanation for the beneficial but also invasive competence of PGPBs, that survive in the host and resist antagonistic effects of the present microbiome, when applied as bio-inoculant ^81^. However, some of the traits also play a role in proliferation within other environments and for pathogenic bacteria, as a consequence of co-evolution to adapt to common environmental conditions ^82^. Examples are siderophore gene clusters, bactericidal compounds, antimicrobial resistance genes, toxin-antitoxins, flagellar systems, pili and secretion systems ^18,83^. Single strains in diverse environments, but also of pathogens, reveal high counts of PGPTs, and might be able to achieve PGP. Such phytostimulatory effects of pathogenic and also human associated strains, like for *Agrobacterium tumefaciens* and *Escherichia coli*, has received attention in the plant-microbe community ^84^. For example, our results confirm that the major known PGP pathway for phosphate solubilization by gluconic acid production and secretion via the *pqq* genes occurs in a number of pathogens, as already discussed in 2013, but detailed phylogenomic analyses are lacking to address their real evolutional dispersion ^85,86^. Further, it is very likely that human and plant virulence factors related to motility, colonization, or effectors might overlap with plant-beneficial traits and do not show significant differences in any group, such as the type VI secretion system (T6SS), known to be important for plant colonization of PGPBs ^87,88^. Finally, it must be considered that bacteria may switch between different lifestyles and, thus, reveal genetic traits for example for both plant symbiosis and pathogenesis, what might hinder a clear separation between both lifestyles during clustering according PGPT traits count only.

We revealed that significant higher genome sizes and gene counts for plant-and soil-associated bacteria correlate with higher PGPT counts, albeit a constant PGPT ratio of approximately one third of all genes present per genome was measured for all environments, which is compatible with the previous study of Levy et al in 2018 ^41^. Our results emphasize much higher genome sizes and gene counts being associated to rhizospheric and / or plant symbiotic strains, rather than by bacteria of other plant spheres and pathogens. While very low coding information is connected to an endo-symbiotic lifestyle, for most of the plant symbionts we cannot conclude the same genomic reduction related to host adaption, as reported for e.g., insect endosymbionts ^89^. Our results suggest such genome reduction events for single strains across all environments, plant spheres and bacteria-induced plant phenotypes but this must be investigated in more detail in the future.

Nevertheless, all results and also the average lower genome sizes of human and animal strains and their pathogens indicate that such an increasing amount of genetic information might be attributed to a higher number of traits related to plant or soil adaption, e.g., plant colonization or higher competitiveness, rather than to general host adaption processes or pathogenicity. Accordingly, recent studies hint to traits related to competitiveness, plant invasion and colonization, as for example evolving antimicrobial and secondary metabolite gene clusters and higher insertion rates of transposases ^90–92^. But our finding about a much higher gene count in plant symbionts, also points to a high number of genes related to the entire plant symbiosis, summarized as PGPTs. That assumption might be also supported by a study of increasing disease-suppressive functions of the endophytic root microbiome apparent after pathogen infection, that detected large, novel biosynthetic gene clusters ^52^

Of course, sole plant growth-promoting efficacy is affected by different aspects of genetic virulent disposition (e.g., virulence factors), genome and domain organization posttranscriptional or posttranslational regulation or the plant genotype, metabolome, microbiome and environmental influences ^40^. In this context, temporal and spatial distribution of symbionts, commensals and their PGPTs in plant microbiomes have to be investigated in more detail, as well, to understand the corresponding variability, even of pathogen invasion efficacy ^77^. Perhaps the abundance of genes and their genomic coordinates might allow one to distinguish between (i) obligate symbionts and / or facultative pathogens, (ii) their host specificity and site adaption processes, rather than using only PGPT occurrences ^93^. We are also aware that not all plant growth-promoting processes might be considered in the current version, as not all genes for all beneficial metabolic pathways are known yet. Here more research must be done, also by *in vivo* experiments. However, any suggestions for new pathways are welcome.

We envision that PLaBAse and the PGPT ontology will help to understand the PGP potential and the microbial community assemblage in its multifaceted manner affected by ecology, plant genotype, plant host immunity, phytochemistry and topology, among others. ^94–96^. Comparable PGPT annotation results will facilitate the selection of plant-beneficial inoculant strains for a sustainable plant production and protection, by determination of (i) best site adapted pathways (e.g., for salinity tolerance) and of (ii) overlapping traits with antagonizing functions of pathogens (e.g., bactericidal compounds), the latter important for exclusion of dysbiosis by competition and assertiveness in an established environment. Those interspecies-interactions, whether symbiotic or not, have to be further elucidated, for example by co-cultivation studies on plant-based media or germ-free plants to determine the best plant-simulating environments ^23,97–99^. Here the cultivability of yet uncultured PA strains will enable a better understanding of genetic traits or genome organization (e.g., biosynthetic gene clusters) related to niche adaption ^100,101^. The prediction of genotypic and phenotypic plasticity of PGPBs could also lead to the selection of promising bio-inoculant candidates from other environments, that are not yet entirely considered ^102^. Thus, our approach should be considered together with the detection of virulence factors (e.g., VFDB ^78^) for biosafety strategies. We further see high potential of our approach in (meta-) genomic, transcriptomic or proteomic analyses (e.g., MEGAN6, gene set enrichment analyses), to elucidate the interkingdom signaling with the plant host under changing climate conditions, to define corresponding synergistic or antagonistic microbial consortia, and to (re-) classify symbiosis islands and plasmids ^3,98,103,104^.

We encourage researchers to contribute to PLaBAse to improve the current tools and to add new features.

## ACKNOWLEDGEMENTS

Within the framework of the study, special thanks go to the support of Andreas Nagel (server & web-domain administration) and to the ZDV of the University Tuebingen, especially to Jens Krueger, who helped us in any issues regarding to HPC on the bwHPC BINAC cluster that is provided by the state of Baden-Wuertemberg (Germany). Further, this work was supported by the German Research Foundation (DFG, INST 37/935-1 FUGG) and by the BMBF-funded de.NBI Cloud within the German Network for Bioinformatics Infrastructure (de.NBI) (031A532B, 031A533A, 031A533B, 031A534A, 031A535A, 031A537A, 031A537B, 031A537C, 031A537D, 031A538A). The Julius Kuehn-Institute - Federal Research Centre for Cultivated Plants in Braunschweig (Germany) must be acknowledged for its contribution to the common hardware deployment. The data analyzed in this research was provided by the Facilities Integrating Collaborations for User Science (FICUS) initiative, the DOE Joint Geno me Institute and the Environmental Molecular Sciences Laboratory, which are DOE Office of Science User Facilities. Both facilities are sponsored by the Office of Biological and Environmental Research and operated under Contract Nos. DE-AC02-05CH11231 (JGI) and DE-AC05-76RL01830 (EMSL). These sequence data were produced by the US Department of Energy Joint Genome Institute http://www.jgi.doe.gov/ in collaboration with the user community.

## AUTHOR CONTRIBUTIONS

Sascha Patz: conceptualization, bioinformatics analysis, programming, writing, reviewing, editing

Anupam Gautam: website programming, writing, reviewing, editing

Matthias Becker: biological evaluation, writing, reviewing, editing

Silke Ruppel: biological evaluation, writing, reviewing, editing

Pablo Rodríguez-Palenzuela: PIFAR tool, reviewing, editing

Daniel H. Huson: conceptualization, writing, reviewing, editing

## ABBREVIATIONS

AA: animal association, animal-associated bacteria
BDSF: *Burkholderia* diffusible signal factor: cis-2-Dodecenoic acid
DB: database
DSF: diffusible signal factor: cis-11-Methyl-2-dodecenoic acid
FA: food association, food associated bacteria
GABA: γ-aminobutyric acid
FLOW: flower(s)
HA: human association, human associated bacteria
IA: industrial association, industrial associated bacteria
ISR: induced systemic resistance
OR: odds ratio
PA: plant-association, plant-associated bacteria
PATH: pathogen
PGP: plant growth-promoting / plant growth-promotion
PGPB: plant growth-promoting bacteria
PGPT: plant growth-promoting trait (gene / protein)
PHYL: phyllosphere
PIFAR: plant interaction and virulence factor resource
PLaBA-db: plant-associated bacteria database
PLaBAse: plant-associated bacteria web resources
Pred: prediction tool
QA: aquatic association, aquatic-associated bacteria
RHIZ: rhizosphere
SA: soil association, soil-associated bacteria
SAR: systemic acquired resistance
SYMB: symbiont
VF: virulence factor
VFDB: virulence factor database

