## Supplemental Figures 1-4 for "PLaBAse: A comprehensive web resource for analyzing the plant growth-promoting potential of plant-associated bacteria"

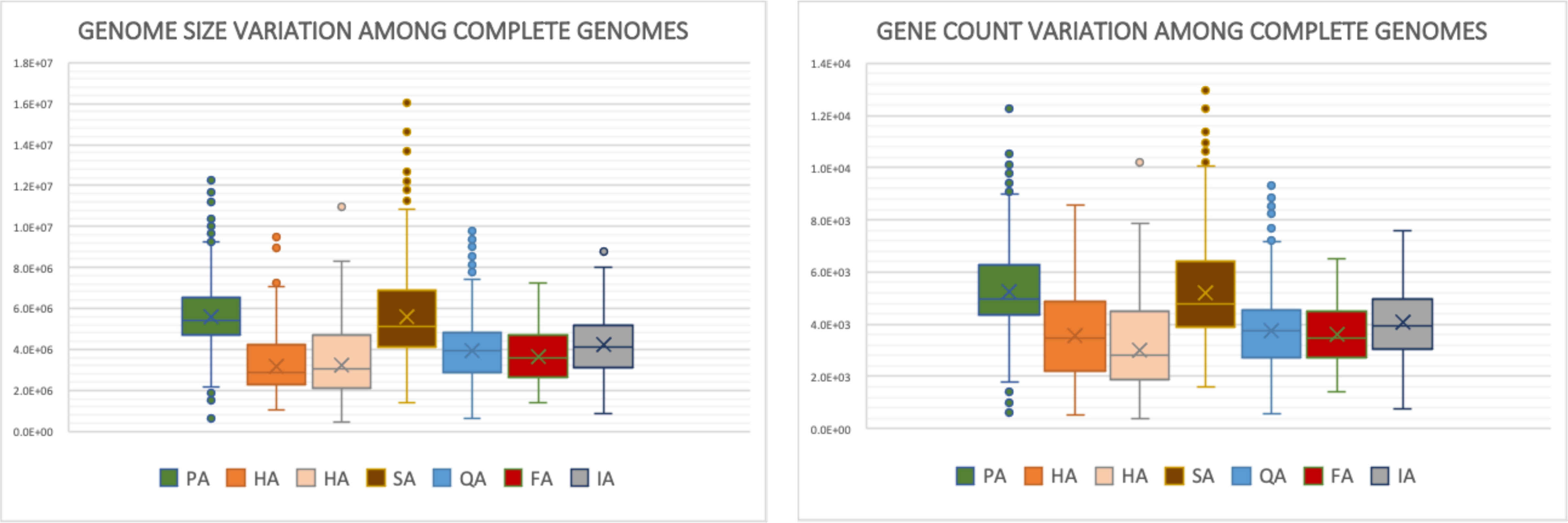


**Supplemental Fig. 1: Genome size and gene count variation among complete genomes only.** Explanations for abbreviations are given in the main paper. The genome size scale has to be understood as basepairs (bp).

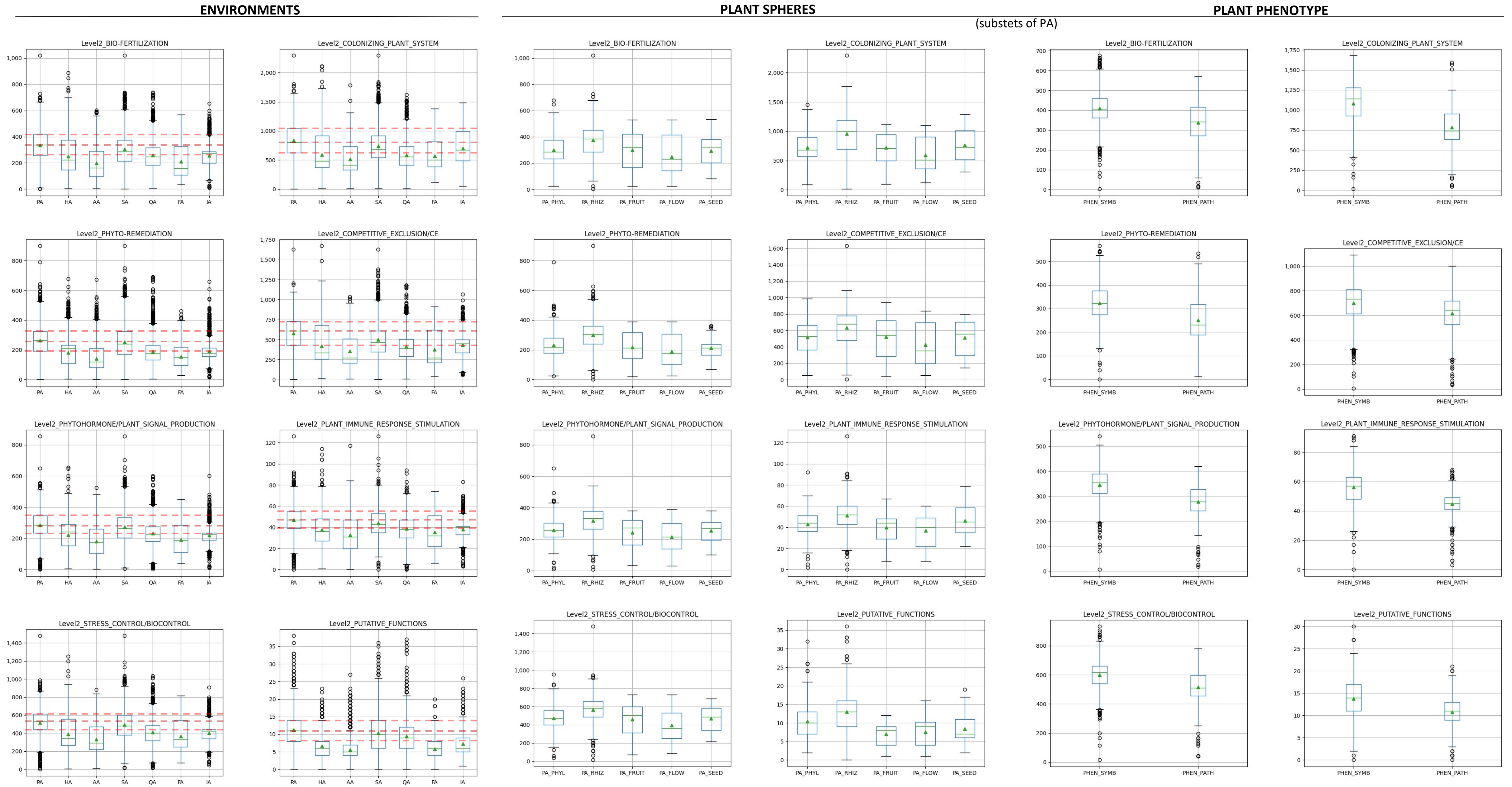


**Supplemental Fig. 2: Variation of PGPT count for all level 2 classes among environments, plant spheres and bacterial plant phenotypes.**
Explanations for abbreviations are given in the main paper.

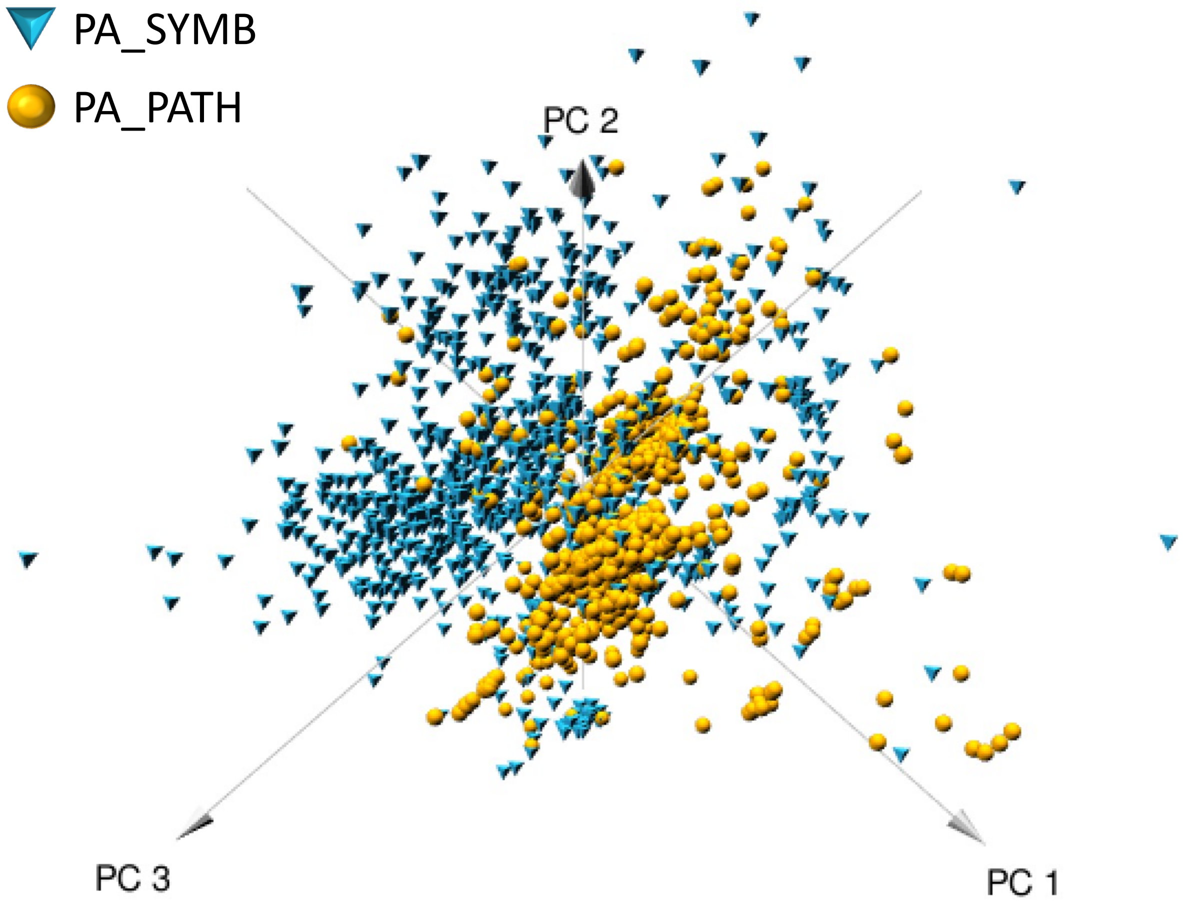


a

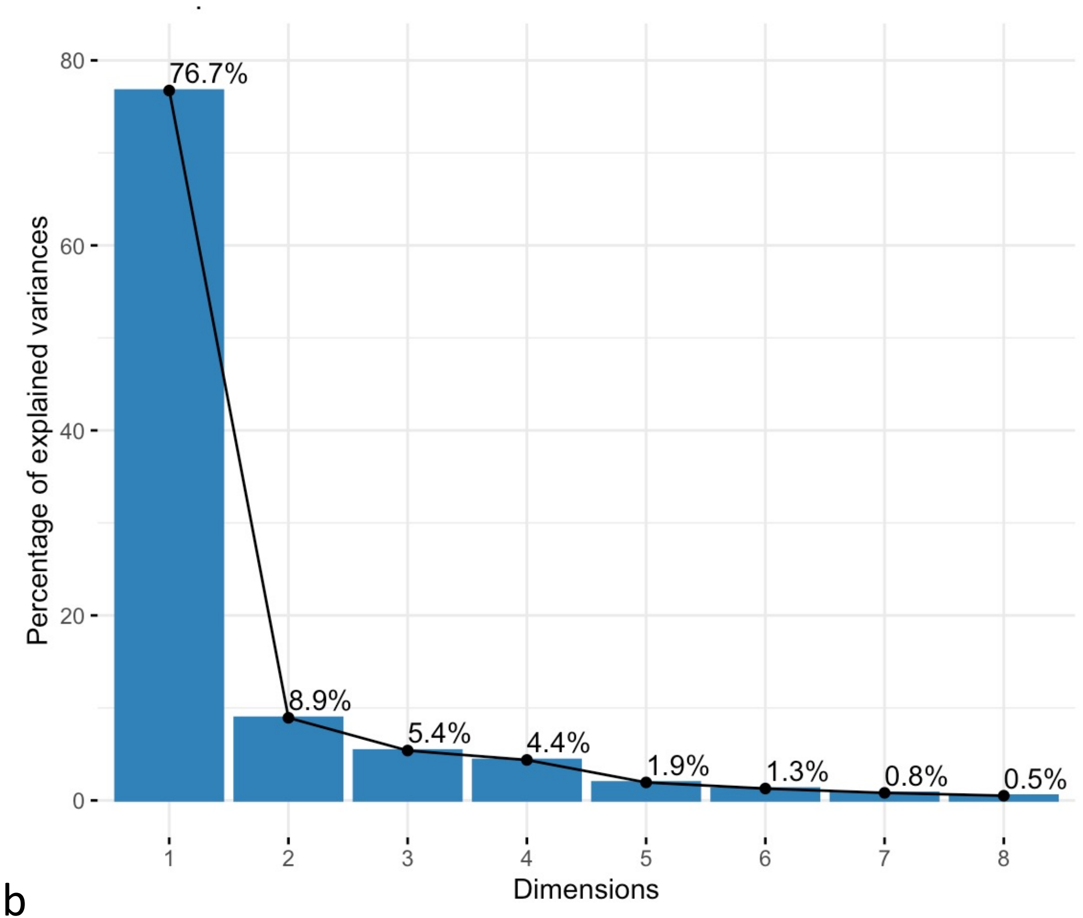


b

**Supplemental Fig. 3: 3D-PCA plot and scree plot for PGPT counts of level 2 classes related to plant symbionts and phytopathogens.** (a) 3D-PCA plot and (b) scree plot of eigenvalues. Explanations for abbreviations are given in the main paper.

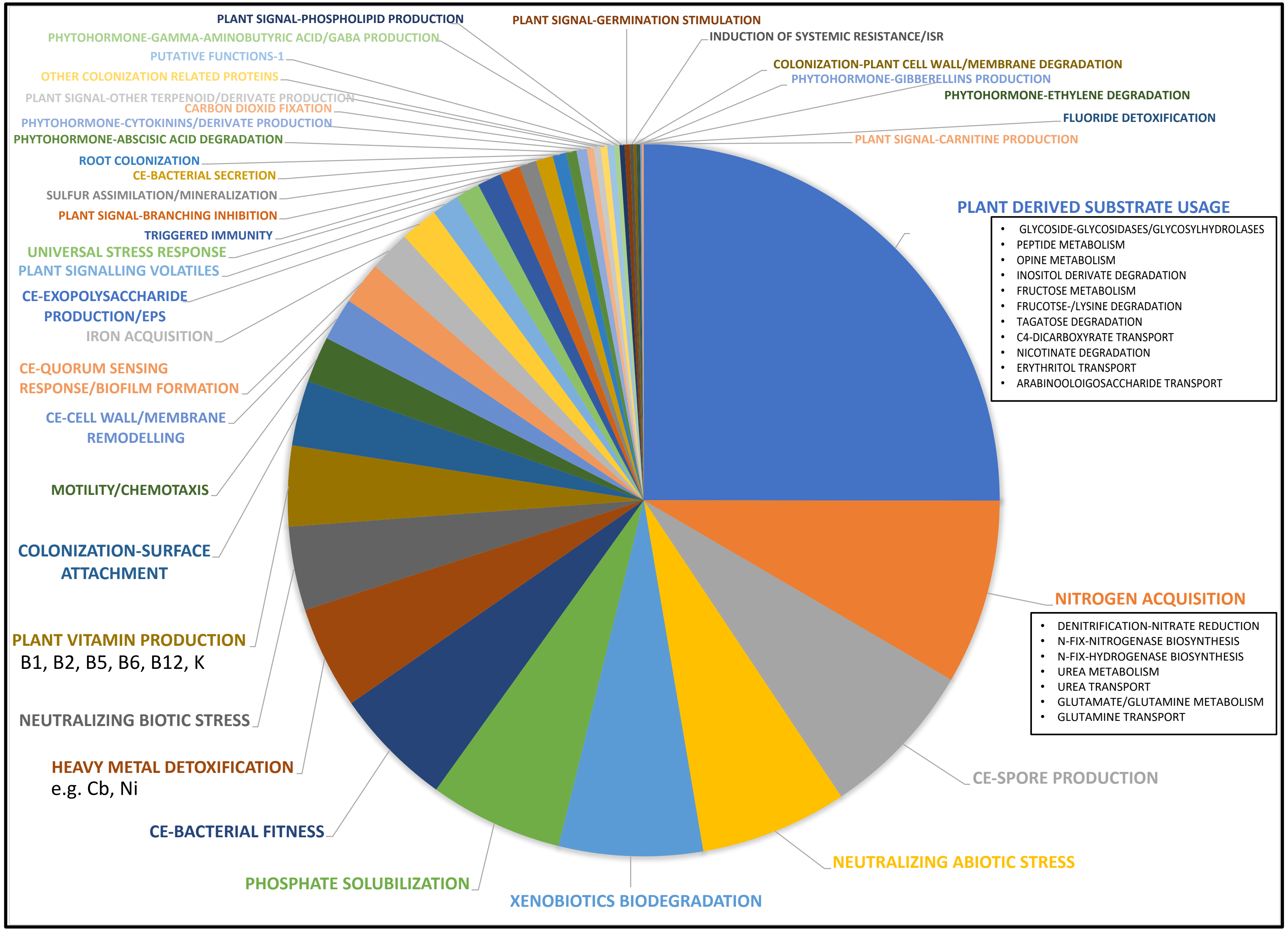


**OTHER EXAMPLES (Level 3 to 5):**

XENOBIOTICS BIODEGRADATION:
- BENZOATE DEGRADATION
- PHENYLACETATE DEGRADATION
- FURFURAL HOMEOSTASIS

P-SOLUBILISATION:
- PYRUVIC ACID BIOSYNTHESIS
- PHOSPHONATE DEGRADATION (*pqq* genes)
- SULFURIC ACID BIOSYNTHESIS

NITROSATIVE/OXIDATIVE STRESS
- ROS REGULATORY PROTEINS

OSMOTIC/SALINITY STRESS:
- GLYCINE-BETAINE METABOLISM
- VITAMIN B3/NIACIN BIOSYNTHESIS
- SODIUM TRANSPORT
- ECTOINE METABOLISM

CE-EXOPOLYSACCHARIDES:
- SUCCINOGLYCAN METABOLIC PATHWAY

COLONIZATION:
- PTI/COLONIZATION-Nod FACTORS
- SURFACE LIPID MODIFICATION
- ADHESINS
- TEICHOIC/COLONIC ACIDS

OTHERS:
- AMINOGLYCOSIDES RESISTANCE-STREPTOM./KANAM./AMIKACIN
- OTHER MULTIDRUG EFFLUX GENES

**Supplemental Fig. 4: Pie chart of enriched PGPTs (level 6) projected back to Level 3 PGPT classes accounting for plant symbiotic strains.**
